# Detecting new allies: Modifier screen identifies a genetic interaction between *Imaginal disc growth factor 3* and a Rho-kinase substrate during dorsal appendage tube formation in *Drosophila*

**DOI:** 10.1101/2020.06.17.156711

**Authors:** Claudia Y. Espinoza, Celeste A. Berg

## Abstract

Biological tube formation underlies organ development, and when disrupted, can cause severe birth defects. To investigate the genetic basis of tubulogenesis, we study the formation of *Drosophila melanogaster* eggshell structures, called dorsal appendages, which are produced by epithelial tubes. Previously we found that precise levels of *Drosophila* Chitinase-like proteins (CLPs), encoded by the *Imaginal disc growth factor (Idgf)* gene family, are needed to regulate dorsal-appendage tube closure and tube migration. To identify factors that act in the *Idgf* pathway, we developed a genetic modifier screen based on the finding that overexpressing *Idgf3* causes dorsal appendage defects with ∼50% frequency. Using a library of partially overlapping heterozygous deficiencies, we scanned chromosome 3L and found regions that enhanced or suppressed the *Idgf3-*overexpression phenotype. Using smaller deletions, RNAi, and mutant alleles, we further mapped five regions and refined the interactions to 58 candidate genes. Importantly, mutant alleles identified *combover (cmb)*, a substrate of Rho-kinase (Rok) and a component of the Planar Cell Polarity (PCP) pathway, as an *Idgf3-*interacting gene: loss of function enhanced while gain of function suppressed the dorsal appendage defects. Since PCP drives cell intercalation in other systems, we asked if *cmb/+* affected cell intercalation in our model, but we found no evidence of its involvement in this step. Instead, we found that loss of *cmb* dominantly enhanced tube defects associated with *Idgf3* overexpression by expanding the apical area of dorsal appendage cells. Apical surface area determines tube volume and shape; in this way, *Idgf3* and *cmb* regulate tube morphology.

## INTRODUCTION

Biological tubes establish the primary design of all organs. For example, the spinal chord is made by the neural tube, which in many vertebrates begins to form by “wrapping” (Nievelstein *et al*. 1993; Catala *et al*. 1996). Wrapping is a mechanism of tube formation in which rows of cells within a sheet constrict their apices while adjacent outer rows of neighboring cells migrate towards each other, zipping together to form a tube that is parallel to the original plane (Lubarsky and Krasnow 2003). Tubes are clearly important both developmentally and physiologically, and yet the signals responsible for inducing epithelial cells to reorganize from a flat sheet into a complex tubular structure are still poorly understood [reviewed by Nikolopoulou *et al*. 2017].

Understanding the genetic programs that drive wrapping is of great interest because their improper implementation gives rise to spinal chord defects such as *spina bifida* and *anacephaly*, which affect ∼ 1 in 1000 births worldwide (Hogan and Kolodziej 2002; Copp *et al*. 2015; Avagliano *et al*. 2019) and represent a major health and economic problem in our society (Bamer *et al*. 2010; Yi *et al*. 2011). Although animal models and GWAS studies have helped identify genetic players that might be involved in tube formation during human development (Copp and De Greene 2010; Wang *et al*. 2019), limited accessibility to the process impedes studies to understand how these genes influence morphogenesis. An important question that remains in the field is exactly how those, and other unknown genetic programs, drive tube formation at a cellular level. Studies in the *Drosophila* system could help us overcome this challenge.

The *Drosophila* laid egg and the egg chamber that produces it serve as great models to study tube development (Berg 2005; Osterfield *et al*. 2017). The laid egg has two eggshell structures, known as dorsal appendages (DAs), which facilitate gas exchange for the developing embryo by retaining pockets of air within the chorion (Hinton 1969). The dorsal appendages develop through the formation of two biological tubes that use the wrapping mechanism of tube formation (Dorman *et al*. 2004) (Figure 1). Although the dorsal appendages are not tubes themselves, cellular tubes mold their physical form (Dorman *et al*. 2004; Berg 2005). Therefore, by looking at the shape of the dorsal appendages on a laid egg, we can assess mistakes during the process of tube formation.

**Figure 1.**
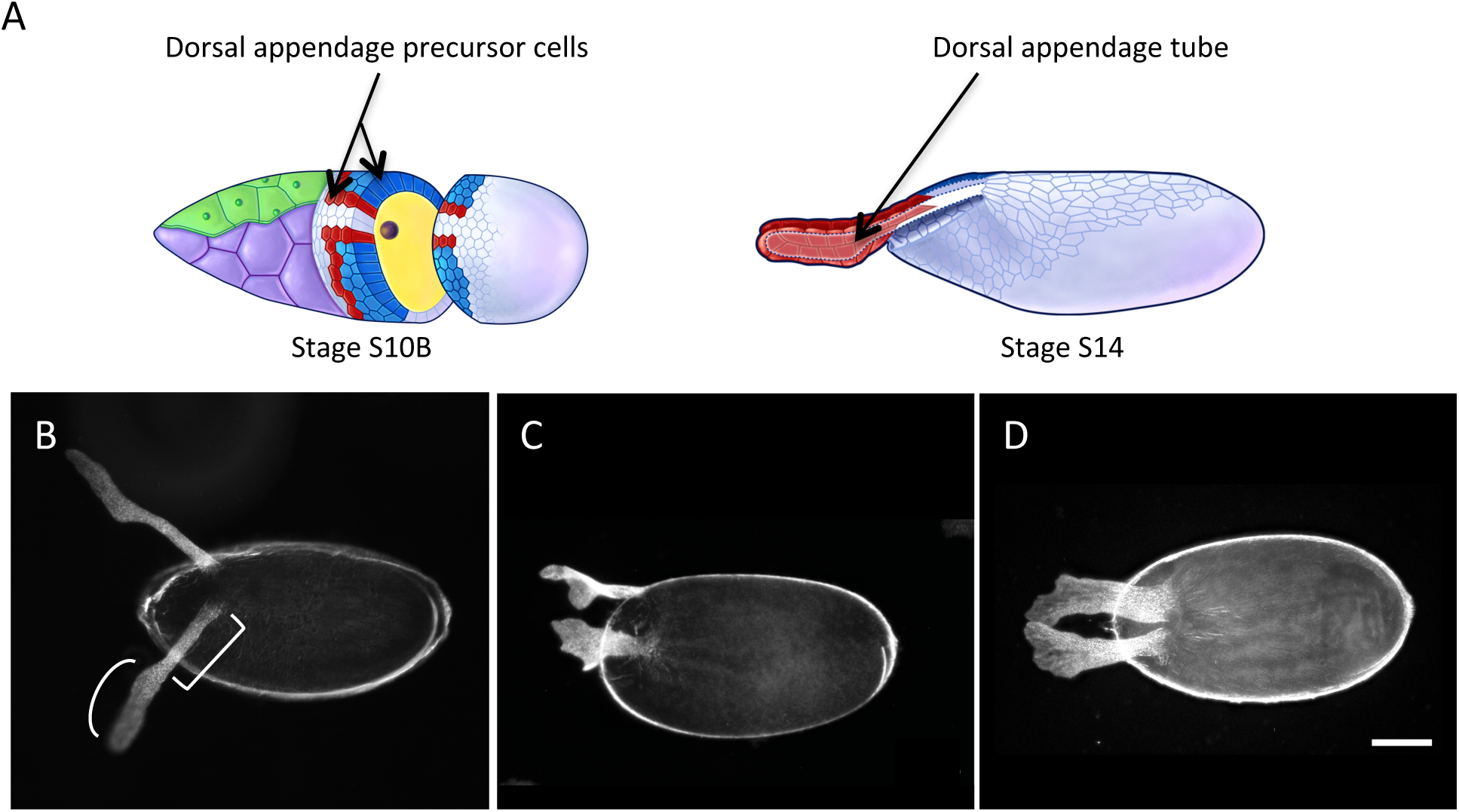
Dorsal appendage formation is a good model to study biological tube formation. (A) Schematic drawings of two later stages of oogenesis, S10B and S14, depicting the cells that make the egg chamber. On the left, arrows indicate the cells programmed to build the dorsal appendages in red (floor cells) and blue (roof cells). On the right, an arrow points to the tube lumen, defined by a dotted line. Drawings obtained with permission from Dorman *et al*. 2004. (B-D) Dark-field micrographs of laid eggs. Anterior is to the left, dorsal is up in B and C, facing out of the page in D. (B) Egg from a wild-type female shows the rounded stalk (square bracket) and flat paddle [curved bracket] of normal dorsal appendages. (C) An egg laid by a *bullwinkle* (*bwk*) loss-of-function mutant exhibits short, wide, dorsal appendages with flat stalks and wavy paddles. (D) An egg laid by an *Idgf3*-overexpression mutant has short, wide, dorsal appendages similar to *bwk*. Pictures in C and D were obtained with permission from Zimmerman *et al*. 2017. The scale bar in D = 100 microns and applies to B, C, and D.

Dorsal appendage formation takes place before the egg is laid, while the egg chamber develops in the ovaries of the adult female (Figure S1A). The egg chamber consists of 16 germ-line cells (a single oocyte connected to 15 sibling nurse cells) surrounded by a mono-layered epithelium of somatic follicle cells (Figure S1B). Within the ovary, the egg chambers are organized in assembly lines, ranging from the youngest, stage-1 (S1) egg chambers, at the anterior, to mature, stage-14 (S14) egg chambers, at the posterior. This organization and the presence of multiple egg chambers within a single female allow us to simultaneously observe tube formation at different stages of development (King, 1970). By dissecting fly ovaries and fixing egg chambers, we can capture stationary processes of otherwise fast-moving events of morphogenesis (Hudson and Cooley 2014; Peters and Berg 2016A).

The morphological changes that form the dorsal appendage tubes begin at S10B of egg chamber development (Figure 1A and Figure S1B). At this stage, the nurse cells occupy the anterior half of the egg chamber and the oocyte occupies the posterior half. The follicle cells that surround the nurse cells have become so thin and squamous they are called “stretch cells”, while the follicle cells over the oocyte are columnar in shape. By S10B, among the columnar follicle cells, two patches of cells are differentially programmed to make the two tubes that mold the dorsal appendages (Figure 1A). From S10B to S14, these cells undergo cell-shape changes, cell intercalation, and cell migration in order to form two mature tubes (Dorman *et al*. 2004; Osterfield *et al*. 2013). While the tubes are forming, the follicle cells release chorion protein into their lumen; these chorion proteins are later cross-linked to create an oar-shaped structure with a rounded stalk and a flat paddle (Fig 1A, B). Once this morphogenetic process is finished, the follicle cells undergo apoptosis and detach from the egg (Nezis *et al*. 2002). Thus, the final shape of the dorsal appendages reflects the process of tube formation (Figure 1B).

We have used the dorsal-appendages model to identify and characterize genes involved in tube formation (Berg 2005). For example, we identified *bullwinkle (bwk)*, which when mutated results in wide and short dorsal appendages resembling moose antlers [hence the name] (Figure 1C; Rittenhouse and Berg 1995). In *bwk* egg chambers, the DA-forming tubes don’t seal properly, and dorsal-appendage-making cells migrate more laterally than anteriorly (Dorman *et al*. 2004). Interestingly, *bwk* acts outside the dorsal-appendage-making cells, in the nurse cells, by regulating expression of signals to the stretch cells (Rittenhouse and Berg 1995). The stretch cells act as mediators to then communicate with the dorsal-appendage-making cells, facilitating the proper formation and elongation of the DA tubes (Tran and Berg 2003). The signals involved in each of these processes are not completely understood.

To understand the molecular landscape of the stretch cells and how it drives dorsal appendage formation, we recently used proteomic analysis to assess stretch cells purified from wild-type and *bwk* egg chambers; we discovered that *bwk* mutants vastly over express a novel family of growth factors, the Imaginal disc growth factors (Idgfs). Lowering the expression of *Idgf4, Idgf5*, or *Idgf6* ameliorates the *bwk* mutant phenotype, suggesting they are part of the *bwk* signaling pathway that regulates dorsal appendage formation. Up-regulating only a single member of this family, e.g., *Idgf3*, is sufficient to produce a phenotype similar to the *bwk* mutant (Figure 1D) (Zimmerman *et al*. 2017).

There is limited knowledge about the *Idgfs*. The first *Idgfs* were identified from conditioned medium and shown to act as growth factors, playing roles in cell-shape changes, cell proliferation, and cell migration in cell lines cultured *in vitro* (Kirkpatrick *et al*. 1995; Kawamura *et al*. 1999). Supporting the hypothesis that Idgfs might influence cell behaviors, transcripts from *Idgf1, Idgf2, Idgf3, Idgf4*, and *Idgf6* accumulate in sites of the embryo where major morphogenetic changes occur, such as the ventral furrow and midgut invaginations (Kawamura *et al*. 1999; Jambor *et al*. 2015). The *Idgfs* encode proteins containing a signal-peptide domain (Zhu *et al*. 2008) and a mutation-bearing-chitinase catalytic domain (Varela *et al*. 2002), suggesting they are secreted molecules that evolved from chitanases but lack the ability to break down chitin. Their receptors and signaling pathway are not known.

In addition to helping us comprehend biological tube formation, elucidating the mechanism of action of the Idgfs could give insight into other human diseases since the human orthologues of the Idgfs, the Chitinase-Like Proteins (CLP’s) (Zhu *et al*. 2008), are up-regulated in immune diseases (reviewed in Ober and Chupp 2009), in numerous cancers (reviewed in Libreros *et al*. 2013), and during infections (Erdman *et al*. 2014). In spite of their well-known association with these diseases, relatively little is known about their mechanism of action. Indeed, it is not clear whether the CLPs are pathogenic, protective, or both, depending on circumstances. Characterizing the cellular mechanisms of Idgfs in tube formation could facilitate studies of human CLPs by providing testable hypothesis on how Chitinase-like proteins could be acting in these contexts.

The main goal of this study was to increase our understanding of the *Idgfs* and their role in tube formation by identifying a genetic pathway that interacts with the *Idgfs* during dorsal appendage formation. We designed an unbiased screen to uncover genes that suppress or enhance the DA defects produced by over-expressing *Idgf3*. We identified large regions of chromosome 3L that, when removed by half, showed a possible genetic interaction with *Idgf3* for tube morphogenesis. Using the same approach with smaller, overlapping deletions, we narrowed down a subset of those possible interacting regions to a few candidate genes. Using RNAi lines and mutant alleles, we discovered a genetic interaction between *Idgf3* and *combover (cmb)*, a Rho-kinase substrate that physically interacts with a Planar Cell Polarity (PCP) pathway component. Through immunostaining, we go on to show how this interaction influences tube formation at a cellular level. In this way, we report the identification and first cellular characterization of an *Idgf-*interacting gene. Additionally, we identify other potential *Idgf3-*interacting candidates, some of which might also interact in the *cmb*— *Idgf3* pathway.

## MATERIALS AND METHODS

### Fly Stocks used

*w*^*1118*^ and the other stocks used in this work are available upon request. *w*^*1118*^ ; *CY2-GAL* (Queenan *et al*. 1997) was provided by Trudi Schüpbach and was used in lieu of a wild type strain. *w*^*1118*^ ; *UAS-Idgf3/TM3,Sb* was obtained from the Bloomington Stock Center (BL# 52658). Using these strains we created the stock *w*^*1118*^; *CY2-GAL4; UAS-Idgf3/TM3, Sb*. All deficiency lines were provided by the Bloomington Stock Center (Table 1). *UAS-RNAi* lines were obtained from the Bloomington or Vienna Stock Centers (Table 2, Ni, *et al*. 2011; Dietzl *et al*. 2007). The *combover* loss-of-function (LOF) strain, *w*^*1118*^ ; *+/+* ; *cmb*^*KO*^*/TM6B, Hu*, and an overexpression allele *w*^*1118*^ *P{UAS-cmb-RB}* (Table 2) were generously donated by Andreas Jenny’s laboratory (Fagan *et al*. 2014). The *cmb*^*KO*^ LOF allele is null for both of the Cmb protein isoforms due to the deletion of an ∼ 1 kb fragment early in the coding region and its replacement with a *white*^+^ marker. The overexpression allele used in this paper produces only the smaller of the two Cmb isoforms, Cmb-PB.

**TABLE 1.**
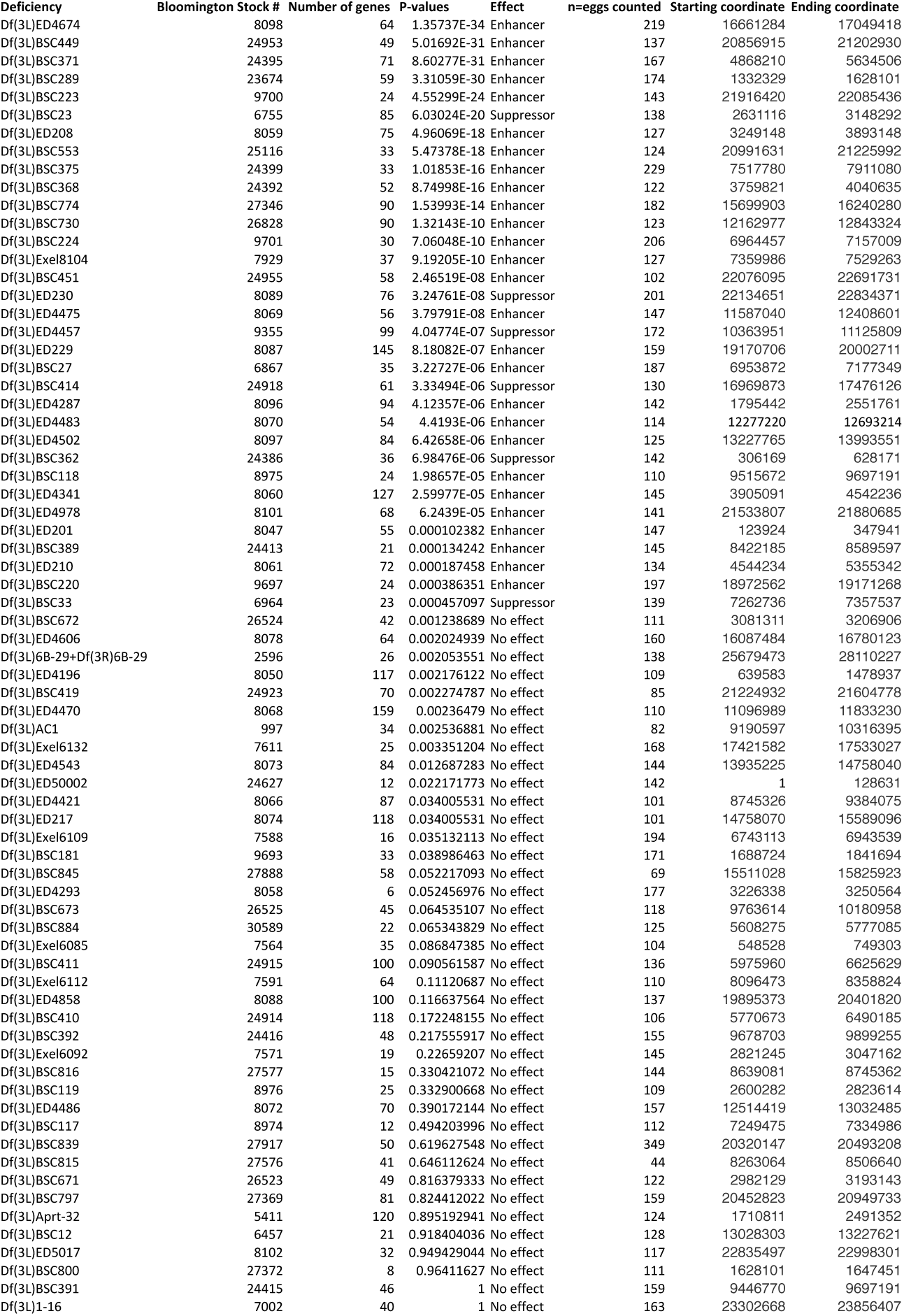

**TABLE 2.**
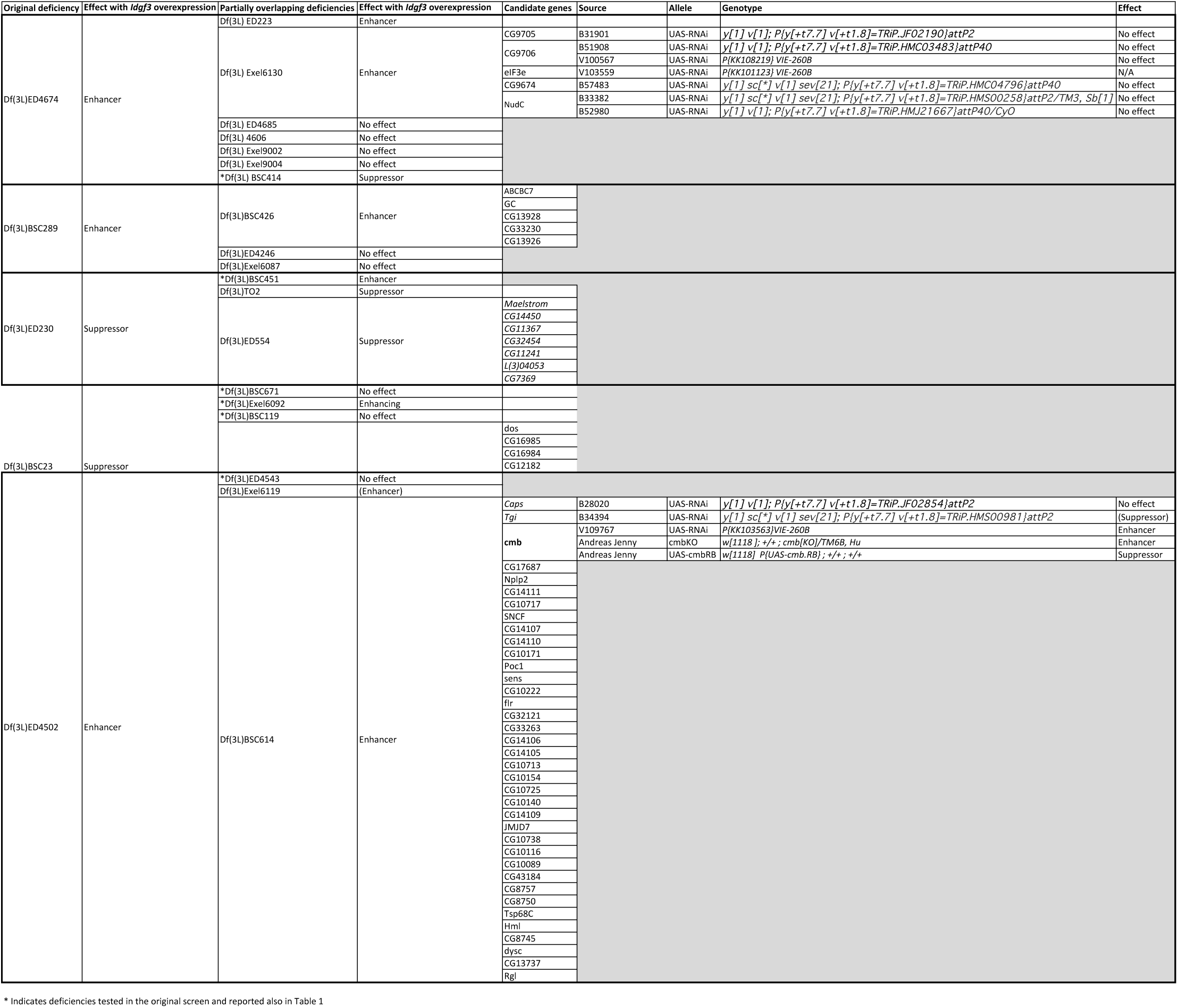

### Modifier-screen crosses

Six-to-ten virgin females from the *Idgf3*-overexpression stock were crossed to four males of each stock from the deficiency kit, or to males from overlapping deletion lines, or to RNAi or mutant-allele strains. For the modifier screen, crosses were done at 25 ^°^C, while for narrowing down regions, crosses were performed at 22 ^°^C. From each cross, at least nine but usually 25 F1 females were used for the egg collection assay (Figure S2).

### Dorsal-appendage analyses

One-day-old to four-day-old females of the desired genotypes were transferred to 30^>°^C in nutrient rich vials with males. Flies were transferred everyday to fresh nutrient-rich vials for three days. On day four, flies were transferred to collection tubes that contained apple juice agar plates with fresh yeast where flies laid eggs. On day five, laid eggs were collected, alternately rinsed with water and embryo wash (0.7% NaCl, 0.05% Triton X-100), mounted on slides in 70 uL of Hoyer’s medium (van der Meer 1977), and incubated overnight at 65°C. Dorsal appendages were scored by using dark-field optics on a Nikon Labophot microscope at 10X magnification; n>100 unless specified in Table 1.

We grouped dorsal appendage phenotypes into three categories (Figure 2A). Eggs with DAs closely resembling wild type were classified as Normal/Mild: the dorsal appendages were positioned just lateral to the dorsal midline, extended anteriorly ∼ 30% of egg length, and exhibited an oar-like shape with paddles that occupied about half the length of the entire dorsal appendage. These DAs had smooth edges. We classified eggs as Moderate when two of the wild-type features looked mildly defective, such as slightly shorter DAs with wavy paddles. Some eggs exhibited stronger defects, such as DAs that were triangular in shape. For others, the DAs were of normal size, but there was no clear separation of the paddles with the bases, as if the paddles were missing. In some instances, the edges of the dorsal appendages looked jagged or serrated. This category also contained proportionally normal-looking dorsal appendages but of increased size relative to the entire egg, which was of normal size. We scored eggs as Severe when the DAs were short (half the normal length) and wide, or when they exhibited defects in three or more of the normal features. In some eggs, the two dorsal appendages were linked by chorion protein in between them. In other instances, there was a small quantity of chorion protein extending out of the egg, but there was not a specific shape. In other eggs, the DAs were merged at their bases, or chorion protein accumulated on the dorsal side of the egg instead of forming DAs.

**Figure 2.**
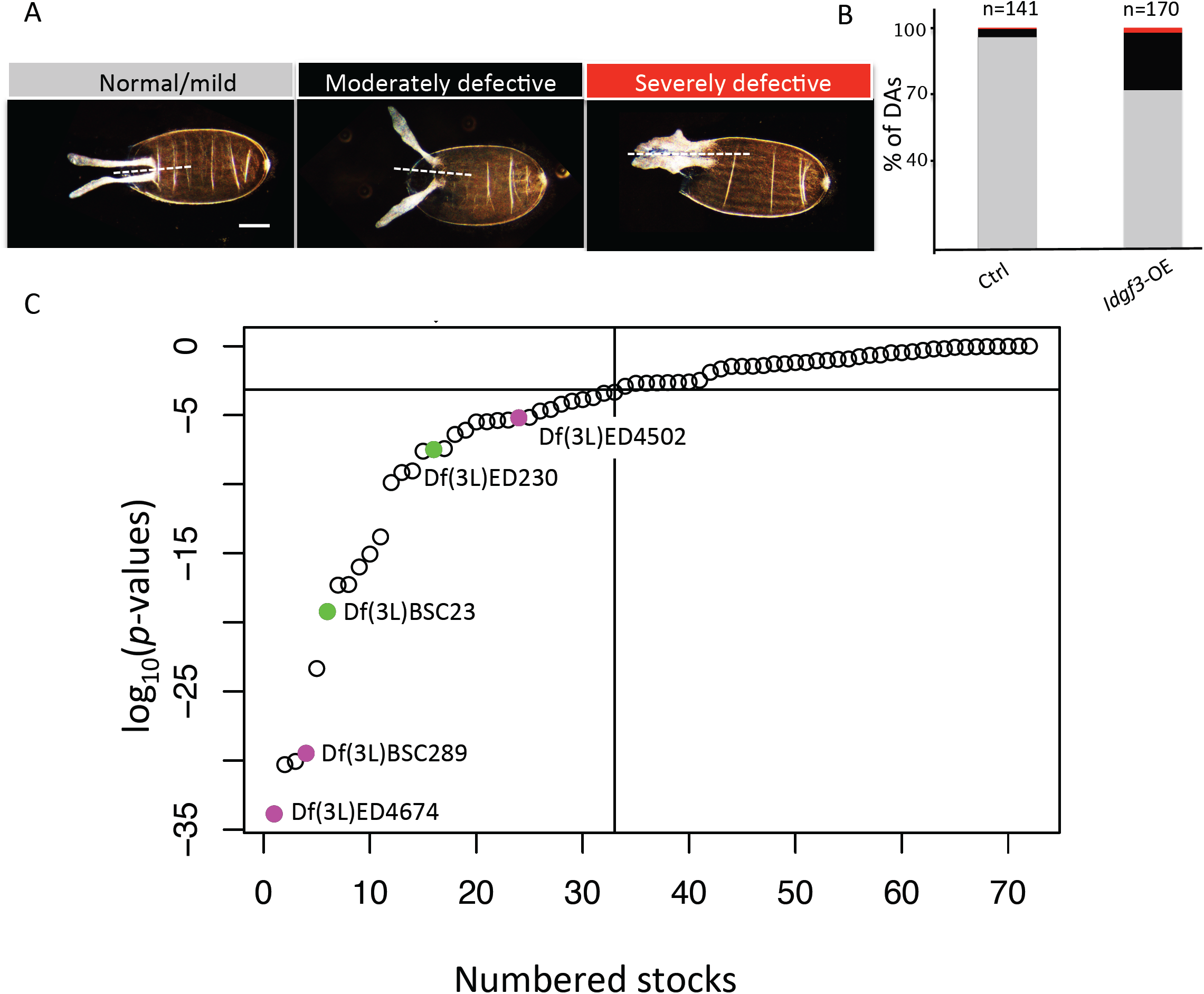
Modifier screen identifies regions on chromosome 3L that enhance or suppress the dorsal-appendage defects due to overexpressing *Idgf3*. (A) Representative images of dorsal appendage phenotypes observed in modifier screen, grouped and color coded into normal/mild (gray), moderately defective (black), or severely defective (red). Images are oriented with anterior facing left and dorsal facing out of the page. Dotted line shows the midline of the egg chamber. Scale bar is 100 microns and applies to all the pictures. (B) Graph: proportions of eggs with normal/mild, moderately defective, and severely defective dorsal appendages produced by females of genotypes: *w*^*1118*^; *CY2-GAL4/+; +/+ (*right) and *w*^*1118*^; *CY2-GAL4/+; UAS-Idgf3/+* (left). (C) log_10_ of Chi-squared *p*-values calculated from comparing the dorsal appendage phenotypes between the *w*^*1118*^; *CY2-GAL4/+;UAS-Idgf3/+* (control) and *CY2-GAL4/+;UAS-Idgf3/Df(3L*) (tested). Significance threshold is y= −3.1, which corresponds to the log_10_ of the Bonferroni correction of 0.05. Filled circles indicate the deletion lines that we chose for further analysis, in magenta (enhancers) and in green (suppressors). Arrows indicate strong and weak interactions.

### Immunostaining

On the third day at 30°C, F1 females from the desired genotypes were anesthetized on a CO_2_ pad and their ovaries dissected. To limit variability between samples, dissections were performed simultaneously by three people in the lab and completed within fifteen minutes. Dissected ovaries were placed in phosphate-buffered saline (PBS) on ice and fixed in 4% EM-grade formaldehyde [Thermo Fisher Scientific, Catalog# 43368] in PBS with 0.1% Tween 20 for twenty minutes. Ovaries were then washed three times in PBS with 0.1% Tween 20. To ensure even staining, single egg chambers of stages S10B and S12 were then dissected out, permeabilized with 1% Triton X-100 in PBS followed by three washes in PBS with 0.1% Tween 20. Eggs were then blocked in 10% Western Blocking Reagent (WBR, Roche) in PBS with 0.1% Tween 20 and incubated with gentle shaking overnight at 4°C with mouse anti-Broad-core (1:250 uL, 25E9.D7 concentrate, Developmental Studies Hybridoma Bank, DHSB; Oda *et al*. 1994) and rat anti-E-cadherin (1:50 uL, DCAD2-concentrate, DHSB; Dubreuil *et al*. 1987). Egg chambers were then washed four times in PBS with 0.1% Tween 20 and 10% WBR and incubated for three hours at room temperature with Alexafluor 488-conjugated goat anti-mouse (1:200), Alexafluor 568-conjugated goat anti-rat (1:200), 4’, 6-Diamidino-2-phenylindole (DAPI) (1 ug/mL) in PBS with 0.1% Tween 20 and 10% WBR. The egg chambers were then washed three times in PBS with 0.1% Tween 20 and 10% WBR, once in PBS 0.1% Tween 20, and mounted in Aqua polymount [Polysciences, Catalog# 18606] for imaging.

### Confocal image acquisition

Imaging and scoring of egg chambers were done blind by covering the genotype labels with tape and assigning letters A, B, C, and D. Their names were revealed after all data analyses. We used the Leica SP8X confocal microscope, with the 20X objective and then with a zoom of _X_2 focusing on the DA-forming patches. Wavelength emissions 488nm, 461nm, and 568nm were used with a PMT detector at 30%, 5% and 30% intensity power, respectively. The format of the acquired images was 1024 x 1024 at a speed of 600, with Z slices separated by 0.25 μm. We captured images moving basally (facing outward in this tissue) to apically (facing the oocyte), starting from where the Br-positive nuclei were visible and ending below the apical E-Cadherin staining, just after reaching the oocyte’s cytoplasm.

### Image analysis

Images were processed using ImageJ Version 2.0.0-rc-59/1.51n (FIJI) (Schindelin *et al*. 2012). To identify DA patches, we created a Z projection using all images captured in the Br-positive channel for each egg chamber. To uniformly distinguish high-Broad staining (DA cells) from low-Broad staining (posterior and lateral cells) among different S10B egg chambers, the images were smoothed and made binary using the method Max Entropy or Momentum. We only used egg chambers in which the entire DA patch was visible, that is, those egg chambers mounted with a dorsal or partially dorsal view of the patches. To measure aspect ratios, we traced the exterior boundary of the high-Br cells to create a shape that enclosed the entire patch. Measurements were set up to calculate shape descriptors: aspect ratio, circularity and roundedness of the basal side of the cells. For apical surface measurements, we used single slices in the E-Cadherin channel at the apical-most region of the tube. We calculated length by tracing and recording a straight line from the base of the dorsal appendage tube to its tip.

### Statistical analysis

For the modifier screen, we used a Chi-squared test for consistency to compare control and test samples. Although we categorized DAs into three groups (normal/mild, moderate, or severe), some samples lacked sufficient severe eggs to conduct the test properly; we therefore combined the moderate and severe groups into a single “defective” class for all comparisons. We used the R statistical package (R Core Team 2017) to generate a list of *p*-values using the function chisq.test() with one degree of freedom. We calculated a threshold of significance using the Bonferroni correction test by dividing the 0.05 significance value by 72 (the number of samples we compared), resulting in a 6.9 _X_ 10^−4^ cutoff. To calculate *p*-values for image analyses, the R function for two-sided unpaired t-testings, t.test(), was used with a confidence interval of 95%.

*Idf3* modifier screen data and antibodies available upon request.

## RESULTS AND DISCUSSION

### Modifier-screen set up

Previous studies from our lab, using the *GAL4-UAS* system, demonstrated that overexpressing *Idgf3* in the stretch cells of the egg chamber causes dorsal appendage defects about 50% of the time (Zimmerman *et al*. 2017). We used this knowledge to identify possible genetic interactions with *Idgf3* by screening regions of the genome that, when reduced to a dosage half that of wild type, could suppress or enhance the frequency of dorsal appendage defects.

We chose chromosome 3L to scan for modifiers of the *Idgf3-*overexpression phenotype because we have limited knowledge of 3L genes that might be involved in dorsal appendage formation (Tran and Berg 2003; Berg 2005; Boyle *et al*. 2010). We used the deficiency kit available at the Bloomington Stock Center (Cook *et al*. 2010) and identified 72 lines that were the minimum number of stocks that uncover virtually all of 3L, a chromosome arm that comprises ∼ one fifth of the genome (Table 1). All the lines have defined breakpoints and share the same genetic background (Parks *et al*. 2004; Ryder *et al*. 2007). Each deletion uncovers on average 58 genes, with a range of six to 159 genes.

We planned to use *UAS-RNAi* constructs to test individual genes within possible interacting regions, but since we did not know which cells of the egg chamber respond to the *Idgf3* signal, we needed a *GAL4* driver that would express in all follicle cells. We created a stock that uses *CY2-GAL4* (Queenan *et al*. 1997) to drive expression of UAS- *Idgf3* in all follicle cells from stage 6 onward (Figure S2B).

We examined the dorsal appendage phenotype of this newly created stock. Flies that expressed Gal4 alone produced eggs with DA defects 4% of the time, while flies that overexpressed *Idgf3* produced eggs with DA defects 29% of the time (Figure 2B). The reduced frequency of DA defects observed with overexpression of *Idgf3* using *CY2-GAL4* compared with *c415*, a stretch-cell-specific *GAL4* driver (50%, Zimmerman *et al*. 2017), could be due to spatial, temporal, or quantitative differences in Gal4 expression (Figure S1C). For example, overexpression of *Idgf3* from the stretch cells could create a signal concentration gradient, while overexpressing *Idgf3* from all the follicle cells could create a uniform signal cloud, exposing the DA-making cells to different amounts of Idgf3. Alternatively, early overexpression of *Idgf3* using *CY2-GAL4* might cause the activation of pathways that counteract the effects of *Idgf3* overexpression, pathways that are ineffective if activated at S10. Although more investigation is needed to discover the mechanism that produces different frequencies of DA defects, the 29% frequency produced by *CY2-GAL4* allowed us to screen for modifiers of the *Idgf3-*overexpression phenotype.

Subsequently, we observed that maintaining the *Idgf3-*overexpression stock at 25°C caused a gradual decline in the frequency of defective DAs over time. Since the *CY2-GAL4* driver is moderately active at 25^>°^C (Queenan *et al*. 1997), we hypothesized that this activity is enough to cause the accumulation of *Idgf3*-overexpression-suppressing mutations. To remedy this problem, we rebuilt the *CY2-GAL4 -> UAS-Idgf3* strain and maintained it at a lower temperature, 22^>°^C. This new strain (when shifted to 30^>°^C as described in the methods), continues to produce eggs with defective DA phenotypes at ∼ 29% frequency.

### Screening for suppressors and enhancers

We crossed females from the *Idgf3-*overexpression stock with males of each of the 72 deficiency lines or with males of a control stock, *w*^*1118*^. By choosing non-Balancer flies, we obtained females that overexpressed *Idgf3* while also being heterozygous for a deficiency uncovering one part of chromosome 3L (Figure S2B); crosses to the control *w*^*1118*^ males produced females that only overexpressed *Idgf3* (Figure S2A). We shifted the flies to 30^>°^C to optimize Gal4 activity, then collected eggs and scored DA defects from each cross.

To determine whether each 3L deletion removed genes that interacted with *Idgf3*, we calculated the frequency of dorsal appendage defects on eggs laid by the *Idgf3*-overexpressing—deletion-heterozygous females and compared that value with the frequency of DA defects on eggs from females overexpressing *Idgf3* alone. We calculated *p*-values as an indicator of strength of interaction (Figure 2C) and drew a threshold of significance at 6.9 x 10^−4^ after correcting for multiple testing (see methods). To our surprise, we found that 46% of the deficiencies significantly modified the *Idgf3-* overexpression phenotype: 38% of the deficiencies enhanced and 8% of the deficiencies suppressed (Table 1).

For this part of the screen, we did not test if each deficiency produced a dorsal appendage phenotype on its own because we wanted to quickly identify possible interacting deficiencies. Therefore, some deletions might have had a dorsal appendage phenotype independent of *Idgf3* overexpression. If so, we over-estimated the number of interacting deficiencies in this first stage of analysis.

We also noticed that our data are skewed towards enhancers. One possible factor contributing to this result is that the Idgfs are dosage sensitive for dorsal appendage formation: both down regulation and up regulation of each *Idgf* cause dorsal appendage defects (Zimmerman *et al*. 2017). Therefore, genes that play a role in either type of regulation could perturb this balance and enhance the frequency of dorsal appendage defects. Also, since each deficiency reduces the copy number of many genes at one time, these large deletions could cause a cumulative effect if more than one cellular pathway is affected. Finally, the dorsal appendages are not critical structures of the fly; we therefore do not expect to find internal mechanisms that ensure proper DA development under such genetic alterations. Nonetheless, in this large screen, we identified several potential sites on chromosome 3L that genetically interact with *Idgf3*.

### Selecting regions to narrow down

We used three criteria to pick deficiencies for further analyses. First, we wanted to identify both suppressors and enhancers of the *Idgf3* overexpression phenotype since these opposite phenotypes could reveal opposing inputs into how Idgfs are regulated or received. Second, we wanted to identify candidate genes that through their interaction with *Idgf3* would reveal critical processes for tube formation. In other words, removing only one copy of the gene while overexpressing *Idgf3* would disrupt tube formation drastically. Those candidate genes would be uncovered by the deficiencies with small *p*-values (Figure 2C, strong intensity). Alternatively, and our third criteria, we were interested in identifying important developmental pathways in which the *Idgfs* might play a role but that have back-up mechanisms to ensure robust function. In other words, removing one copy of a gene will produce only a mild effect on the *Idgf3*-overexpression phenotype because redundant genes or genes in parallel pathways could make up for the reduction of the gene product. Those genes would be uncovered by deficiencies with *p*-values near the threshold of significance (Figure 2C, weak intensity).

These three criteria led us to choose *Df(3L)ED4674, Df(3L)BSC289, Df(3L)BSC23, Df(3L)ED230* and *Df(3L)ED4502* for further study (Figure 2C, Table 2).

### Narrowing down the interacting regions

We narrowed down the regions of interaction by using overlapping deficiencies. First, we took advantage of a tiling effect obtained from the original screen. 72% of the deficiencies from the primary screen partially overlapped with other deletions adjacent to their ends (Table 1, overlapping coordinates). Additionally, we obtained smaller, partially overlapping deficiencies from the Bloomington Stock Center, deficiencies that were produced using the same technologies as those used in the original modifier screen (See examples in Figure 3A and 3B).

**Figure 3:**
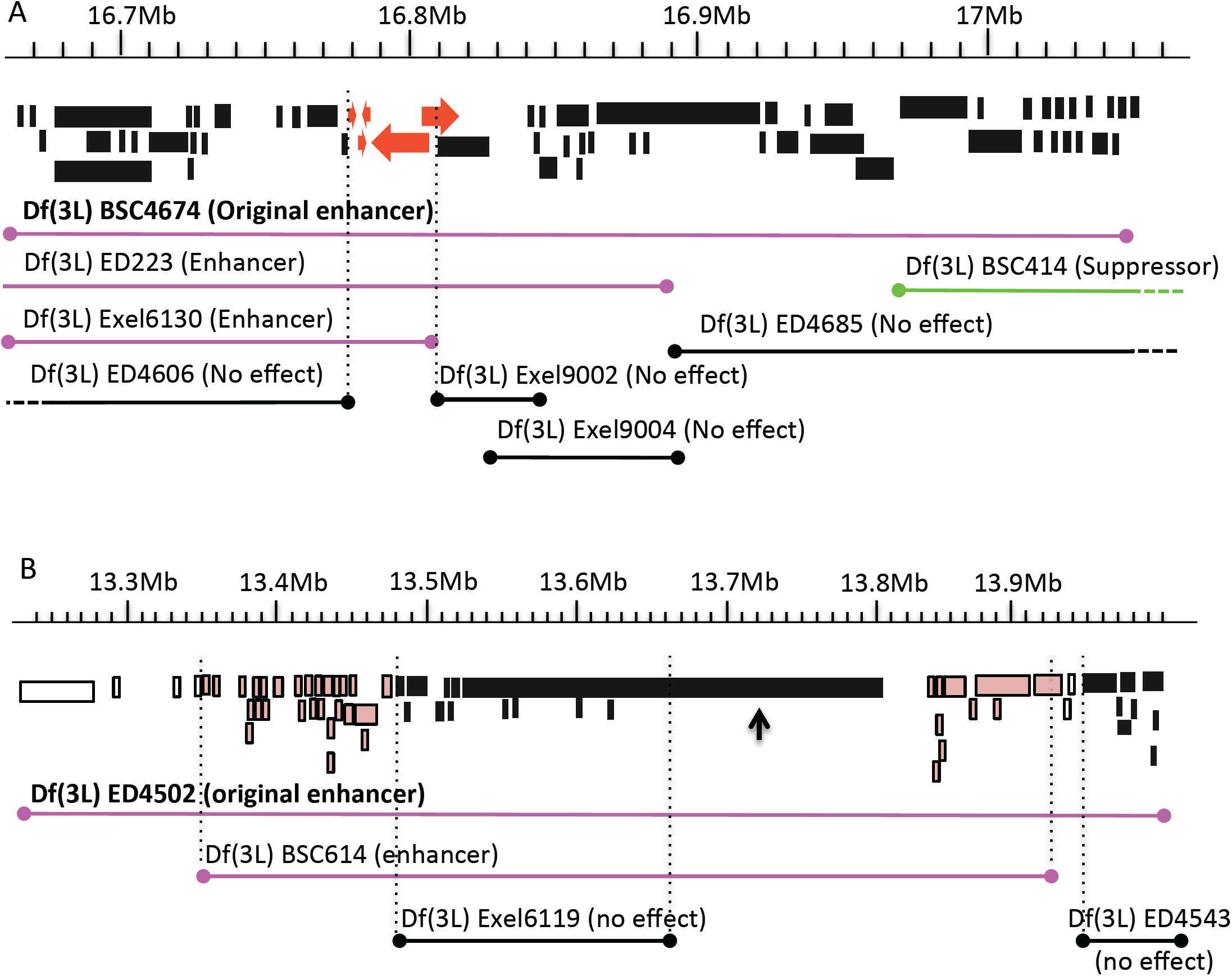
Overlapping deficiencies across chromosome 3L were used to narrow down the regions that interact with the *Idgf3* gene. (A, B) Genome browser maps showing two examples of interacting regions and the deletions we used to refine the list of potential interacting genes. Each block indicates the coding and non-coding regions of a gene. Unfilled blocks are genes whose interaction was neither confirmed nor discarded; black blocks are genes that failed to interact with *Idgf3*; red or pink blocks indicate potential candidate genes. Magenta lines show the span of the deficiencies that enhanced the *Idgf3*-overexpression phenotype. A green line shows the span of a deficiency that suppressed the *Idgf3*-overexpression phenotype. Black lines show the span of the deficiencies that did not change the percentage of defects observed by overexpressing *Idgf3*. Dotted regions indicate deletions that extend distally or proximally from the region shown here. (A) Example of a simple narrowing-down process. *Df(3L)BSC4674* strongly enhanced *Idgf3* overexpression. *Df(3L)ED223* and *Df(3L)Exel6130* confirmed and narrowed down the region of enhancement; these deletions did not produce a phenotype on their own. All the other deficiencies tested in the region did not have any effect on the *Idgf3*-overexpression phenotype, narrowing down the region to five candidate genes, shown in red. (B) Example of a complex narrowing-down process. *Df(3L)ED4502* identified an enhancer region, which was confirmed and narrowed down with *Df(3L)BSC614*. Two other deficiencies uncovered portions of the region; neither produced any effect on the *Idgf3*-overexpression phenotype. These results left 37 candidate genes remaining to be tested. The small arrow points to a gene that was discarded as a candidate gene because the coding regions for all its transcripts lay within the non-interacting deficiency. Ruler above each browser map shows the span of DNA in Megabases (Mb).

Our strongest enhancer, *Df(3L)BSC4674*, overlapped with another deficiency from the original modifier screen, a deletion that suppressed the *Idgf3-*overexpression phenotype (Figure 3A). This result eliminated those genes in the proximal region of *Df(3L)BSC4674*. We tested six additional overlapping deficiencies in the region and found two that strongly enhanced the *Idgf3*-overexpression phenotype. Although these deletions increased the frequency of DA defects in the *Idgf3*-overexpression background, they did not result in a phenotype when *Idgf3* levels were normal (not shown), demonstrating that the defects depended on overexpressing *Idgf3*. By assessing the regions of overlap among these deletions, we ascertained the presence of an *Idgf3-*interacting gene among five candidate genes.

To narrow down the interacting regions uncovered by *Df(3L)BSC289, Df(3L)ED23*, and *Df(3L)ED230*, we used a similar approach and identified 5, 4, and 7 candidate genes, respectively (Table 2).

When investigating *Df(3L)ED4502*, we found one overlapping deficiency, *Df(3L)BSC614*, that enhanced the *Idgf3-*overexpression phenotype, confirming a genetic interaction with *Idgf3* and narrowing down the region to 51 candidate genes (Figure 3B). To refine the interaction further, we tested two additional deficiencies available in the region and found they did not enhance the *Idgf3-*overexpression phenotype. One of those deficiencies, *Df(3L)Exel6119*, resulted in a high number of dorsal appendage defects when *Idgf3* levels were normal, suggesting that a gene in that region plays a role in dorsal appendage formation but does not interact with the *Idgf3* pathway. Since this small deficiency is contained within the original deficiency *Df(3L)ED4502*, it explains why *Df(3L)ED4502* also results in dorsal appendage defects when *Idgf3* is at normal levels (Figure 4, lane 3). In contrast to *Df(3L)Exel6119*, however, *Df(3L)ED4502* interacted synergistically with *Idgf3*. We concluded that the genes uncovered by *Df(3L)BSC614*, but that fall outside *Df(3L)Exel6119*, were responsible for the enhancing effect seen with the large deficiency *Df(3L)ED4502* in an *Idgf3* overexpression background (Figure 4, lane 4). In total, our mapping suggested 37 candidate genes.

**Figure 4:**
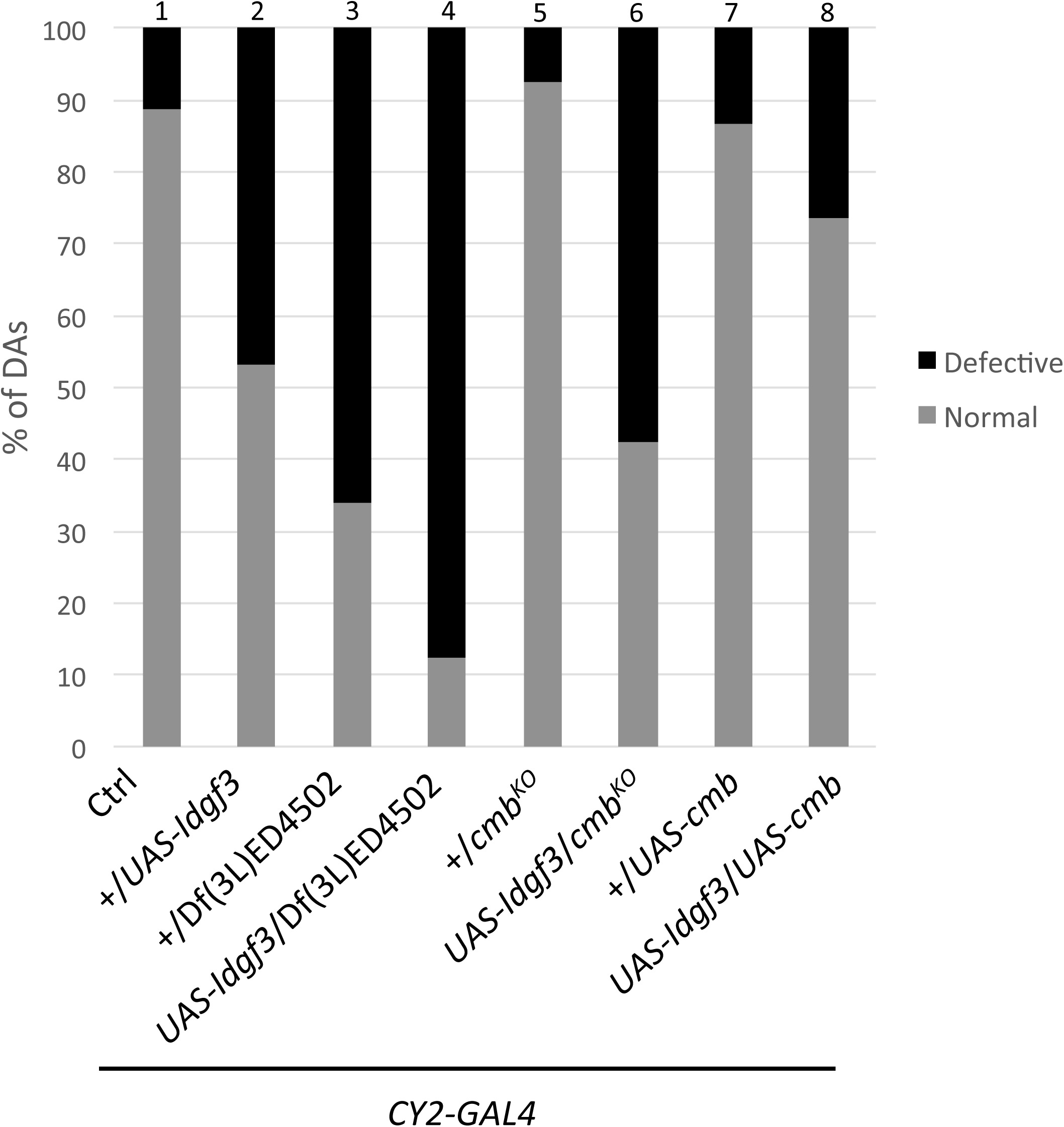
*cmb* genetically interacts with *Idgf3*. Dorsal appendage phenotypes were quantified as normal (gray) or defective (black). Control female *(w*^*1118*^; *CY2-GAL4/+)* produced approximately 11% eggs with defective dorsal appendages (lane 1). The *Idgf3*-overexpression females (*w*^*1118*^; *CY2-GAL4/+;UAS-Idgf3/+)* produced approximately 46% of eggs with defective dorsal appendages (lane 2). *Df(3L)ED4502* uncovered the *cmb* gene and produced a phenotype that was enhanced by the overexpression of *Idgf3* (lane 3 and lane 4). Removing one copy of *cmb* resulted in a small number of defective dorsal appendages (lane 5), similar to the *CY28GAL4* control (lane 1). Removing one copy of *cmb* in an *Idgf3*-overexpression background (lane 6) enhanced the *Idgf3*-overexpression phenotype (lane 2). Overexpressing *cmb* resulted in a small number of defective dorsal appendages (late 7). Overexpressing *cmb* simultaneously with *Idgf3* (lane 8) suppressed the *Idgf38* overexpression phenotype.

### Identifying interacting genes

To identify the actual gene responsible for the interaction with *Idgf3*, we used *UAS-RNAi* constructs generously provided by the Bloomington or Vienna Stock Centers. Using the same DA assay, we tested those alleles for interaction with *Idgf3*. When evaluating the *UAS-RNAi* lines, we kept in mind that the amount of reduction of the gene product would likely differ from the original screen, in part because knockdown depends on the amount of Gal4 protein, which also drives expression of *Idgf3*. Since we were reducing transcripts only in the follicle cells, and the germ line produces a large amount of RNA to be loaded maternally into the embryo, we could not assess the level of transcript reduction easily. Furthermore, protein products could perdure even if transcripts were completely degraded. We therefore interpreted a lack of phenotype from RNAi knockdown with caution, and we used null alleles when available, as nulls more closely mimic the conditions of the original deletion screen.

We narrowed down the region uncovered by *Df(3L)ED4674*, our strongest enhancer, to five candidate genes (Figure 3A, Table 2). RNAi alleles against four candidates, *CG9705, CG9706, CG9674*, and *nudC*, did not modify the *Idgf3-* overexpression phenotype. When we attempted to knock down expression of the fifth gene, *eIF3e*, by crossing a *UAS-RNAi* construct to either the *Idgf3-*overexpression stock or *CY2-GAL4* alone, pupae did not hatch and we were unable to obtain adult females to test their eggs for dorsal appendage defects. We also failed to obtain adults when using a temperature-sensitive *gal80* with *CY2-GAL4*. Unfortunately, no mutant alleles exist. Our observations suggest that *eIF3e*, which encodes a translation initiation factor, is an important developmental gene in *Drosophila*. Consistent with this hypothesis, the mammalian *eIF3e* is essential for mouse embryonic development (Sadato *et al*. 2018). As more null alleles become available, we will be able to test *eIF3e*, and other individual genes, for interaction with *Idgf3* with more confidence.

### *Idgf3* genetically interacts with *combover*

After refining the interacting region for *Df(3L)ED4502*, one of our modest enhancers, 37 genes remained to be tested (Figure 3B, Table 2). We used a candidate gene approach and tested alleles of three genes whose function is relevant for development (Table 2). We identified a mild genetic interaction with a *combover* (*cmb)* RNAi line, which enhanced the *Idgf3-*overexpression phenotype. We tested a loss-of-function mutant and an overexpression line for this gene (Fagan *et al*. 2014) and found complementary effects: *+/cmb*^*KO*^ enhanced the *Idgf3*-overexpression phenotype, and the overexpression of *cmb-RB* suppressed the *Idgf3*-overexpression phenotype in three separate replicas (Figure 4).

We then asked: how are *Idgf3* and *cmb* interacting at the cellular level to impact the shape of the DA tubes? We tested two hypotheses: 1) they modify the narrowing and lengthening of the tube by regulating cell intercalation; and 2) they control the surface area of the tube lumen.

### *Idgf3* and *cmb* do not affect cell intercalation during dorsal appendage formation

The gene *cmb* was first identified as a substrate of Rho kinase, *in vitro* (Fagan *et al*. 2014). This report suggested that *cmb* might play a role in the Planar Cell Polarity pathway (PCP) since the overexpression of *cmb* in the *Drosophila* wing causes the growth of multiple wing hairs, a phenotype that is characteristic of mutations in other PCP components. Moreover, Cmb physically interacts with one component of the PCP pathway (Multiple wing hairs) in both a yeast two-hybrid system and by immuno-precipitation (Fagan *et al*. 2014).

PCP genes set up planar directionality of the epithelium, defining the orientation of static tissues such as those that produce mammalian hairs, bird feathers, and fish scales (reviewed by Butler and Wallingford 2017). The PCP pathway also coordinates the behavior of cells in morphogenetic tissues, directing movements that drive cell intercalation during a variety of developmental processes (Keller *et al*. 2000; Wallingford and Harland 2001; Park and Moon 2001; Darken *et al*. 2002).

In our system, cell intercalation facilitates tube formation (wrapping) at stage 11, and it helps narrow and lengthen the tubes during S12 – S13 (Dorman *et al*. 2004; Osterfield *et al*. 2013; Ward and Berg 2005). When the DA patches are defined at S10B, they are longer along the Dorso-Ventral (DV) axis compared to the Anterior-Posterior (AP) axis (Figure 5A, I). During S11, the floor cells zip up the tube underneath the roof cells, and the roof cells contract their apices. At the same time, more lateral cells move toward the dorsal midline, exchanging neighbors and altering the shape of the DA patch (Dorman *et al*. 2004; Osterfield *et al*. 2013; Ward and Berg 2005). During S12, as the roof cells release apical tension in a biased fashion (Peters and Berg 2016B), cell intercalation continues, producing a patch that is now longer along the AP axis than the DV axis (Figure 5E, J; Dorman *et al*. 2004; Ward and Berg 2005).

**Figure 5:**
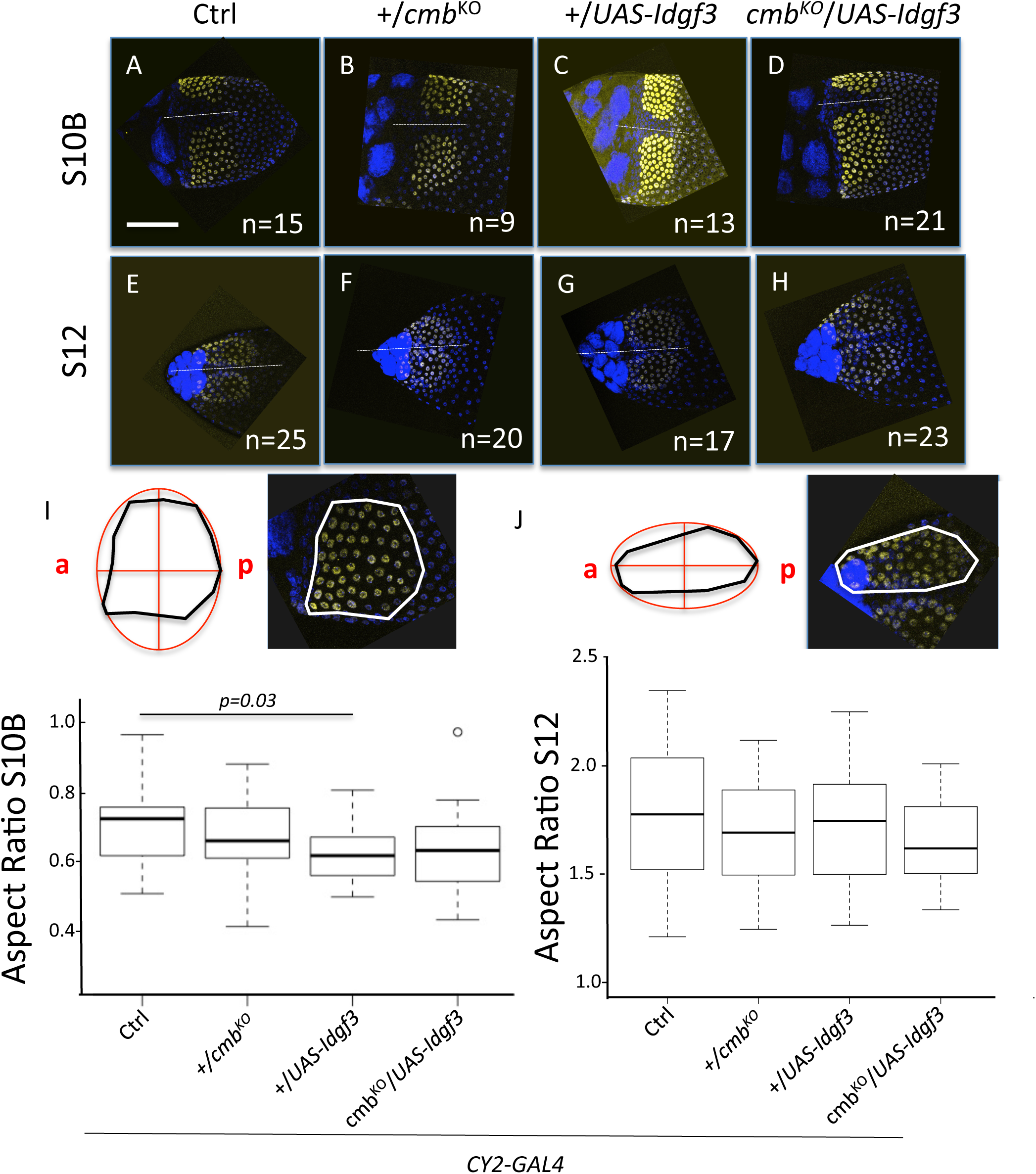
*Idgf3*-*cmb* genetic interaction does not affect cell intercalation during dorsal appendage tube formation. (AIH) Representative examples of central dorsal views of S10B egg chambers (AID), and anterior dorsal views of S12 egg chambers (E-H). Scale bar is 100 microns and applies to all pictures. (A and E) *w*^*1118*^; *+/CY2-GAL4*. (B and F) *w*^*1118*^; *+/CY2-GAL4; +/cmb*^*KO*^. (C and G) *w*^*1118*^; *+/CY2-GAL4;+/ UAS-Idgf3*. (D and H) *w*^*1118*^; *+/CY2-GAL4; UAS-Idgf3/cmb*^*KO*^. Each image is a projection of confocal slices (average ∼38) showing the dorsal appendage patches stained for Broad protein (roof cell nuclei, yellow), and DAPI (DNA, blue). The number of egg chambers scored per genotype is indicated on each panel. Dotted lines show the midlines of each egg chamber. (I, J) Quantification of the aspect ratio of each dorsal appendage patch. Schematics on each panel show the outline area of each dorsal appendage patch, enlarged to the right, that was considered for aspect-ratio calculation using FIJI (See methods). “a” indicates anterior, “p” indicates posterior. Box plots show the mean, quartile, and range of aspect ratios measured for each genotype.

To test if the *Idgf3-cmb* interaction was affecting cell intercalation during DA formation, we quantified this transition by measuring the aspect ratio of S10B and S12 DA patches. We compared egg chambers produced by *CY2-GAL4* (control) females with those produced by *+/cmb*^*KO*^ females, *Idgf3-*overexpressing females, and *cmb*^*KO*^ */Idgf3*-overexpressing females. If *Idgf3* and *cmb* affected cell intercalation, we expected to see similar aspect ratios among genotypes at S10B, but significant differences between control and experimental groups at S12. To mark the exact boundary of the dorsal appendage patches, we used an antibody against Broad (Br), a transcription factor required to specify DA-forming cells (Tzolovsky *et al*. 1999): high levels of Broad (“High Br”) define the DA patches, while moderate or low Broad (“low Br”) marks lateral and posterior main-body follicle cells (Dorman *et al*. 2004). To avoid introducing any bias, we conducted the image acquisition and quantification blind to the genotypes we were analyzing (see methods).

High-Br staining revealed the basal location of the nuclei in the dorsal-appendage-making cells (Figure 5A-H). As expected, at S10B, the shapes of the patches were similar for all genotypes (Figure 5A-D), except for a small but significant difference between the *Idgf3-*overexpression group and the control. Since the means of the aspect ratios of all the genotypes were tightly clustered (0.66+/-0.04; Figure 5I), however, this small difference (*p* = 0.03) might simply have resulted from slight timing differences in S10B stage.

When looking at S12 egg chambers, we were surprised that the DA patches from both control and experimental egg chambers had elongated and narrowed to a comparable degree (Figure 5E-H), with aspect ratios that again exhibited similar means (1.71+/-0.06; Figure 5J). Moreover, the slight differences among genotypes at each stage were not significant (P > 0.05), and the small difference seen in the *Idgf3-* overexpression group at S10B was not present at S12. Based on these results, we concluded that the *Idgf3-cmb* interaction was not directly affecting cell intercalation in our system. This result was consistent with studies in the *Drosophila* testis demonstrating a non-PCP role for *cmb* in sperm individualization (Steinhauer *et al*. 2019).

### *Idgf3* and *cmb* affect apical area of dorsal appendage tubes

To examine the effect of the *Idgf3-cmb* interaction on tube lumen morphology, we stained egg chambers with an antibody against E-Cadherin (E-Cad) to reveal cell shapes (Figure 6A-D). E-Cad localizes on the apico-lateral sides of cells and is an important component of cell junctions, controlling cellular adhesion (Ratheesh *et al*. 2012).

**Figure 6:**
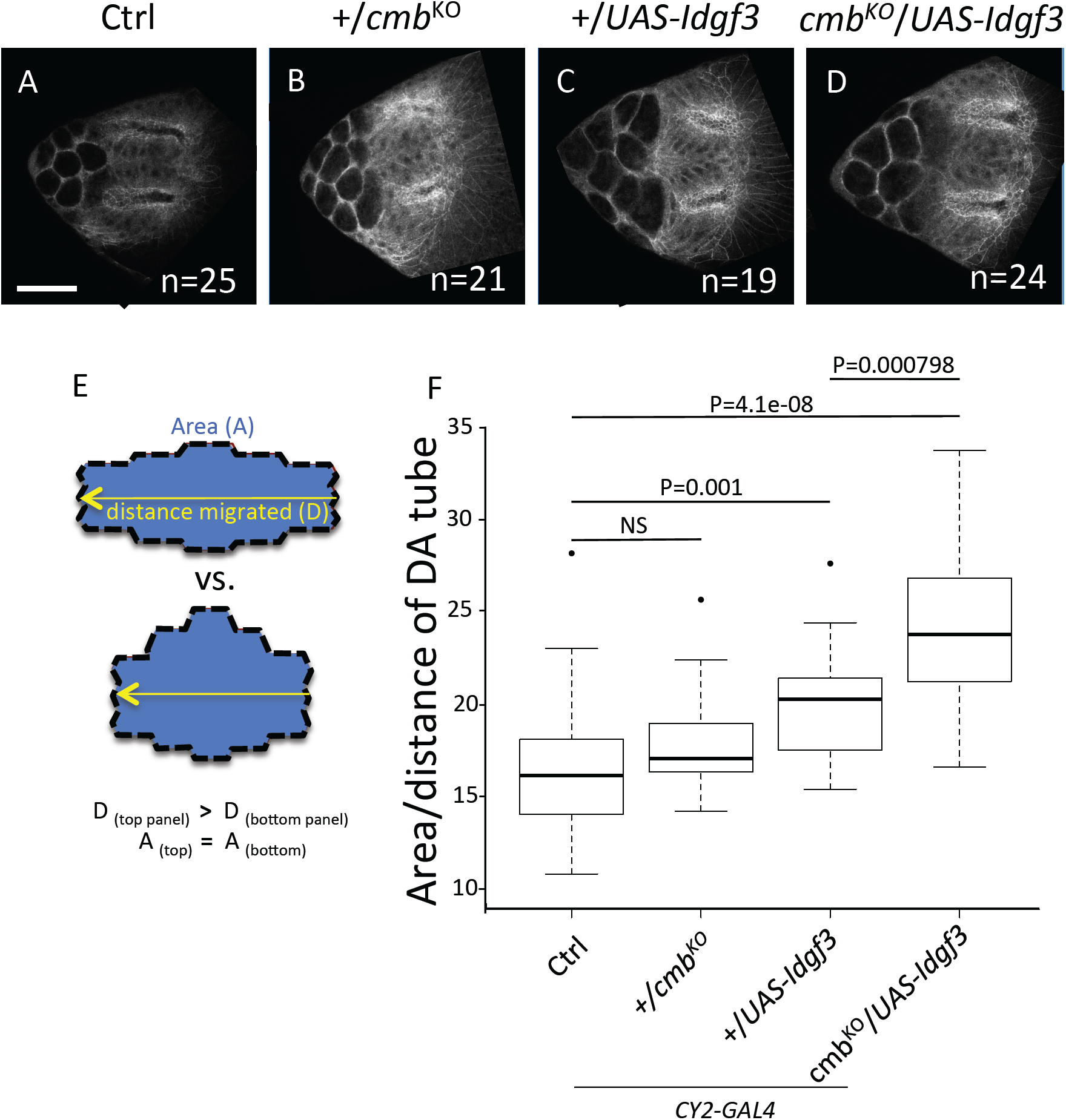
*Idgf3-cmb* genetic interaction affects apical area of the dorsal appendage tubes. (AID) Representative examples of anterior dorsal views of S12 egg chambers stained for E-Cadherin protein. Scale bar is 50 microns and applies to all pictures. (A) *w*^*1118*^; *+/CY2-GAL4*. (B)) *w*^*1118*^; *+/CY2-GAL4; +/ cmb*^*KO*^. (C) *w*^*1118*^; *+/CY2-GAL4; +/UAS-Idgf3*. (D) *w*^*1118*^; *+/CY2-GAL4; UAS-Idgf3/cmb*^*KO*^. Images are a single, 0.5-micron slice taken at the most apical region of the cells that make the dorsal appendage tube. The number of egg chambers scored for each genotype is indicated on each panel. (E) A schematic drawing showing two factors that affect apical tube measurements: i) The temporal progression of dorsal appendage tube elongation (yellow arrows), since cells normally expand their apical surfaces as they move anteriorly and ii) Apical area of tube-making cells (blue space). F) Quantification of the area of the tube normalized by tube elongation (See methods). Box plots show the mean, quartile, and range of aspect ratio/distance migrated measured for each genotype.

We quantified the area of the apical side of the dorsal appendage tubes at stage S12 (Figure 6). Importantly, we had to account for the migration of the dorsal appendage tubes because tube migration, with its accompanying cell rearrangements and apical surface expansion, is an ongoing process during stage S12 (Peters and Berg 2016B). Slight differences in the distance migrated during this stage of DA development could affect apical area (Figure 6E). To account for this problem, we normalized the area of the apical side of the dorsal appendage tubes by the length of the tubes (see methods).

We found that the apical area of the dorsal appendage tube, when controlled by the distance migrated at S12, averaged 16.50+/-1.5 μm in eggs laid by control flies (Figure 6A, 6F). The normalized apical area was similar in eggs laid by +/*cmb*^*KO*^ mutants (18.04+/-1.2 μm; Figure 6B, 6F). In contrast, eggs laid by the *Idg3-* overexpression females had significantly higher values (20.045 +/-1.4 μm; Figure 6C, 6F) than the controls, and knocking out one copy of *cmb* in the *Idgf3*-overexpression background significantly enhanced the *Idgf3*-overexpression effect (24.20 +/-1.7 μm). Since *cmb*^*KO*^*/+* did not produce a phenotype on its own, this change in apical area was dependent on the overexpression of *Idgf3*, similar to what we found in the genetic analysis of laid eggs.

### Role of *Idgf3* and *cmb* during dorsal appendage formation

To explain how *cmb* and *Idgf3* might interact together to regulate the apical area of cells during DA tube elongation, we propose two mechanisms. Both mechanisms rely on the fact that *cmb* is a substrate for Rho kinase (Rok) and that Rok directly affects actomyosin network tension (Munjal *et al*. 2015; Riento and Ridley 2003). Changes in actomyosin network tension likely control the behavior of DA-making cells during tubulogenesis.

During wrapping, in the first steps of tube formation, roof cells constrict their apices and floor cells elongate to seal the tubes. These cell behaviors require high apical network tension (Osterfield *et. al* 2013). Following this step, tubes begin to elongate. Transcription factors that modulate expression of actin-regulatory genes control this transition, and changes in tension then allow biased apical expansion and anterior crawling (French *et al*. 2003; Boyle *et al*. 2010; Peters *et al*., 2013). Since *cmb* is a Rok substrate, it is possible that *cmb* might be involved in regulating tension in dorsal-appendage-making cells during this transition. Experimental work measuring the actomyosin network tension in a *cmb* mutant could help test this hypothesis.

Under this hypothesis, how can *Idgf3* interact with *cmb* to regulate dorsal-appendage-tube elongation? One idea is that *Idgf3* acts upstream of the *Rok-cmb* pathway. That is, *Idgf3* could interact with an unidentified receptor in the dorsal-appendage-making cells, activating Rok and thus inhibiting *cmb* and reducing actin polymerization, as seen in wing hair formation (Fagan *et al*. 2014).

To explain our experimental observations, removing only one copy of *cmb* might not be enough to cause changes in actin polymerization on its own, but overexpressing *Idgf3* might be enough to push the system over a threshold. In this way, the combination of *Idgf3*-overexpression and the removal of one copy of *cmb* might enhance the effect of *Idgf3-*overexpression alone. Analyzing the amounts and distribution of actin in the dorsal-appendage-making cells of *Idgf3*-overexpression and *cmb*^KO/+^/*Idgf3-*overexpression mutants could provide evidence to support or reject this hypothesis.

There is another mechanism that can explain our observations. While *cmb* regulates apical tension of the dorsal appendage tube, *Idgf3* could be acting in a parallel pathway to *cmb*. This alternative mechanism recognizes that apical expansion is normally coordinated with anterior crawling (Boyle and Berg 2009), which involves roof cells physically interacting with the extracellular matrix (ECM) (Dorman *et al*. 2004). We propose that *Idgf3* influences this interaction by modulating the stiffness of the ECM (Zimmerman *et al*., 2017), possibly by activating enzymes that degrade the ECM around the elongating dorsal appendage tubes. Since the leading cells contact the ECM along their basal surfaces, lowering the stiffness of the ECM could lower the resistance the leading cells encounter and alter integrin-mediated intracellular actin dynamics. These changes would then trigger the coordinated expansion of apical surfaces regulated by *cmb*. A similar change in ECM stiffness is critical for the cell-shape changes that drive *Drosophila* wing elongation (Diaz-de-la-Loza *et al*. 2018).

In our experimental observations, removing one copy of *cmb* might not be sufficient to expand apical area because the ECM exerts a force containing the elongating tubes. Overexpressing *Idgf3*, which leads to an abnormal decrease of the ECM stiffness around the tubes, could lower the ECM force that counteracts the expansion of the tubes. Under these circumstances, removing one copy of *cmb* enhances cell expansion, resulting in the phenotype seen in the *cmb*^*KO/+*^; *Idgf3* mutants. Experiments in which we artificially manipulate the stiffness of the ECM could help us understand if the ECM does play a role in dorsal appendage formation. In addition, quantifying the width of the ECM in *Idgf3* and *cmb* mutants would provide evidence to support or reject our hypothesis.

Although more work is needed to fully understand the *cmb*-*Idgf3* interaction, we have successfully identified one biological effect on tube shape when the expression of these genes is altered. It would be interesting to explore if human CLPs affect apical cell area of an epithelium or if the human orthologue of *combover, PCM1*, contributes to actin dynamics in metastatic cancer when the CLPs are up regulated (Libreros *et al*. 2013).

## ACKNOWLEDGEMENTS

Thanks to Andreas Jenny for providing us with the *cmb* alleles; to Lindsay Charley for identifying the method for patch recognition of the S10B egg chambers; to Ken Jean-Baptiste for statistics discussions; to the Berg lab for helpful discussions and input on experiments; to the Bloomington and Vienna Stock Centers for providing stocks; and to the Developmental Studies Hybridoma Bank for antibodies. This work was supported by a NSF Predoctoral Fellowship # 14-590 to CE and NIH R01 GM079433 to CAB.

